# Heat shock protein gene expression is higher and more variable with thermal stress and mutation accumulation in *Daphnia*

**DOI:** 10.1101/2021.01.05.425442

**Authors:** Henry Scheffer, Jeremy Coate, Eddie K. H. Ho, Sarah Schaack

## Abstract

Understanding the genetic architecture of the stress response and its ability to evolve in response to different stressors requires an integrative approach. Here we quantify gene expression changes in response to two stressors associated with global climate change and habitat loss—heat shock and mutation accumulation. We measure expression levels for two Heat Shock Proteins (HSP90 and HSP60)—members of an important family of conserved molecular chaperones that have been shown to play numerous roles in the cell. While HSP90 assists with protein folding, stabilization, and degradation throughout the cell, HSP60 primarily localizes to the mitochondria and mediates *de novo* folding and stress-induced refolding of proteins. We perform these assays in *Daphnia magna* originally collected from multiple genotypes and populations along a latitudinal gradient, which differ in their annual mean, maximum, and range of temperatures. We find significant differences in overall expression between loci (10-fold), in response to thermal stress (~6x increase) and with mutation accumulation (~4x increase). Importantly, stressors interact synergistically to increase gene expression levels when more than one is applied (increasing, on average, >20x). While there is no evidence for differences among the three populations assayed, individual genotypes vary considerably in HSP90 expression. Overall, our results support previous proposals that HSP90 may act as an important buffer against not only heat, but also mutation, and expands this hypothesis to include another member of the gene family acting in a different domain of the cell.

## Introduction

Members of the heat shock protein (HSP) gene family perform an array of essential functions including: acting as molecular chaperones, facilitating the immune response, regulating apoptosis, and signaling protein degradation (Höhfeld et al., 2001; Queitsch et al., 2002; Czarnecka et al., 2006; Javid et al., 2007). The HSP family was first discovered (Ritossa, 1962) and described in *Drosophila melanogaster* (Tissiéres et al., 1974), but has since been the object of intense study across kingdoms and domains (Gupta 1995, Carra et al. 2017). Although HSPs have long been known to act as molecular chaperones aiding in both *de novo* folding and refolding of proteins (Feder & Hofmann, 1999), they also interact with proteins in numerous other contexts (e.g., to facilitate ligand binding or assembly of multiprotein complexes). Interestingly, HSP expression, and the general heat shock response (HSR), is mounted not only in response to heat, but also to a variety of other stressors (e.g., heavy metals, oxidative stress, cytotoxic agents, and mutation; Neuhaus-Steinmetz et al., 1997; Kim et al., 2014; Liu et al., 2015, Queitsch et al., 2002).

Here, we assess the influence of both thermal stress and mutation accumulation on expression levels of two heat shock proteins (Heat Shock Protein 90 (HSP90) and 60 (HSP60)), as well as assessing variation among genotypes and populations in this response. HSP90 is a 90 kDa chaperonin, known as ‘central modulator' or a ‘hub of hubs' due to its role in signaling pathways and protein-protein interactions (Schopf et al. 2017, Zabinsky et al., 2019b), that stabilizes a large clientele of intracellular proteins and signaling proteins. HSP60 is a 60 kDa chaperonin primarily localized to the mitochondria (Cheng et al., 1989). It is involved in the *de novo* folding and refolding of imported proteins in the mitochondria (Martin et al., 1992). HSP60 has also been found in the cytosol where it can participate in either promoting or inhibiting apoptosis (Chandra et al., 2007).

Understanding how organisms respond to thermal stress is an area of urgent biological interest given the current projections of anthropogenically-induced climate change. Variation in HSP expression in response to thermal stress has been demonstrated in a variety of systems (e.g., Tomanek and Somero 2002, reviewed in Feder & Hofmann, 1999). Intraspecific variation in expression profiles within and among populations has not been as widely explored (but see review by Favatier et al. 1997). Among populations, genes thought to respond to heat have been examined in the genus *Fundulus* and individuals vary in their response depending on whether they originated from the Northern or Southern hemisphere, where water temperatures differ (Picard & Schulte, 2004). In addition, the activation of the HSR has been linked to the acclimation of an individual to a given thermal environment, which might explain differences between populations and individuals within a population (Buckley & Hofmann, 2002). While intraspecific variation is posited to be important for resilience to global climate change (Des Roches et al., 2018, 2020), long term thermal tolerance may be attributed to changes in gene expression rather than sequence differences in protein-coding regions (e.g., in corals; Palumbi et al., 2014) raising the question of how acclimation facilitates microevolutionary change (Pauwels et al., 2007, Gienapp et al., 2008).

The role of HSPs as buffers against mutation was initially proposed over 20 years ago (Rutherford and Lindquist, 1998) and has been demonstrated in both animal and plant systems (Queitsch et al., 2002). Because missense mutations can promote protein misfolding and HSPs aid in correct folding, HSP90 has gained a reputation as a “capacitor for mutation” by providing an additional barrier between genotype and phenotype (Jarosz & Lindquist, 2010). The idea is that buffering against protein misfolding stores variation that can then be ‘released' if the cellular pool of HSP90 becomes depleted (Jarosz et al., 2010), as has been demonstrated by mutant lines, knockouts/knockdowns of HSP90, pharmacological interference, and among natural populations that vary in their HSP90 expression (Rohner et al., 2013, Hummel et al., 2017, Mason et al., 2018). A mutation accumulation experiment with hypermutator strains of yeast revealed an enrichment of HSP90 expression (Zabinsky et al., 2019a), underscoring the need for a deeper understanding of the impact of mutation and of intraspecific variation in patterns of HSP expression. There is evidence that upregulation of the bacterial homolog to HSP60, GroEL, can buffer mutations in a similar capacity to HSP90 (Sabater-Muñoz et al., 2015), however it is still unknown if HSP60 buffers mutations in mitochondrial proteins.

We quantify HSP90 and HSP60 expression changes in response to heat shock and mutation accumulation (MA) among different genotypes and populations of *Daphnia magna*. *Daphnia* (Cladocera) have served as an ecological, evolutionary, and ecotoxicological model for well over a century (Schaack, 2008, Shaw et al., 2008, Yampolsky et al., 2014), and genomic resources are now available as well (e.g., Colbourne et al., 2011, Orsini et al., 2016, and Lee et al., 2019). Previously, the *Daphnia* system has been used to demonstrate differences in gene expression, protein production, and evidence for microevolutionary change at HSP genes in the lab in response to environmental change (Pauwels et al. 2007, Mikulski et al., 2009, 2011, Becker et al., 2018). We predict both heat shock and mutation accumulation will increase HSP expression for both genes if they both act as mutational capacitors, but also that the interaction of the two stressors might have a synergistic effect on expression levels, compared to one stress alone. Furthermore, we predicted baseline expression levels and/or the response to stress might differ among populations, but not among genotypes, given the regional differences between Finland, Germany, and Israel. Our experimental design allows us to measure HSP expression levels along multiple axes of comparison, and thus quantify responses to extrinsic and intrinsic stress (heat shock and mutation accumulation) as well as natural variation in basal and stress-induced HSP expression (e.g., among genotypes or populations). Assessing the levels of gene expression variation for HSPs along multiple treatment axes is an important first step towards elucidating the possible role of HSPs as cellular buffers or mutational capacitors and has implications for understanding the evolution of stress responses across lineages and over time.

## Materials and Methods

### Study System and Experimental Design

*Daphnia magna* are aquatic microcrustaceans (Order: Cladocera) with a cosmopolitan distribution that can reproduce quickly, with or without sex. The individuals used in this study were derived from genotypes originally collected in Finland, Germany, and Israel (provided by D. Ebert in 2014), from populations selected because of the distinctive environmental regimes they experience (including temperature, periods of dry down, and census population sizes; Lange et al. 2015) along a latitudinal gradient (see Supplemental Table S1a). In this experiment, we assayed one genotype from Finland (FC), one genotype from Germany (GA), and three genotypes from a single population in Israel (IA, IB, and IC; Figure 1). For the genotype from Germany (GC) and one of the genotypes from Israel (IA), both descendants of the originally collected genotypes (referred to as ‘control lines' hereon) and descendants of five mutation accumulation (MA) lines initiated from each of these clones in 2013 (average number of generations of mutation accumulation = 24; see Ho et al. 2019 for MA details) were assayed (Table S1b). This design allowed us to assess gene expression differences between genes, with and without heat shock, among populations (Finland, Germany, and Israel), among genotypes within a population (within Israel), and between lineages with and without mutation accumulation (Figure 1). Individuals were reared concurrently for 15 days in June and July 2019 in Percival environmental chambers under strictly controlled laboratory conditions to assess levels of heat shock protein (HSP60 and HSP90) expression. Although we set up 4 biological replicates for each lineage/condition combination, in some cases individuals did not survive until the end of the experimental period. In most cases, we were able to perform the RNA extractions and downstream molecular analyses on 2-3 biological replicated for each lineage/condition assayed.

**Figure 1.**
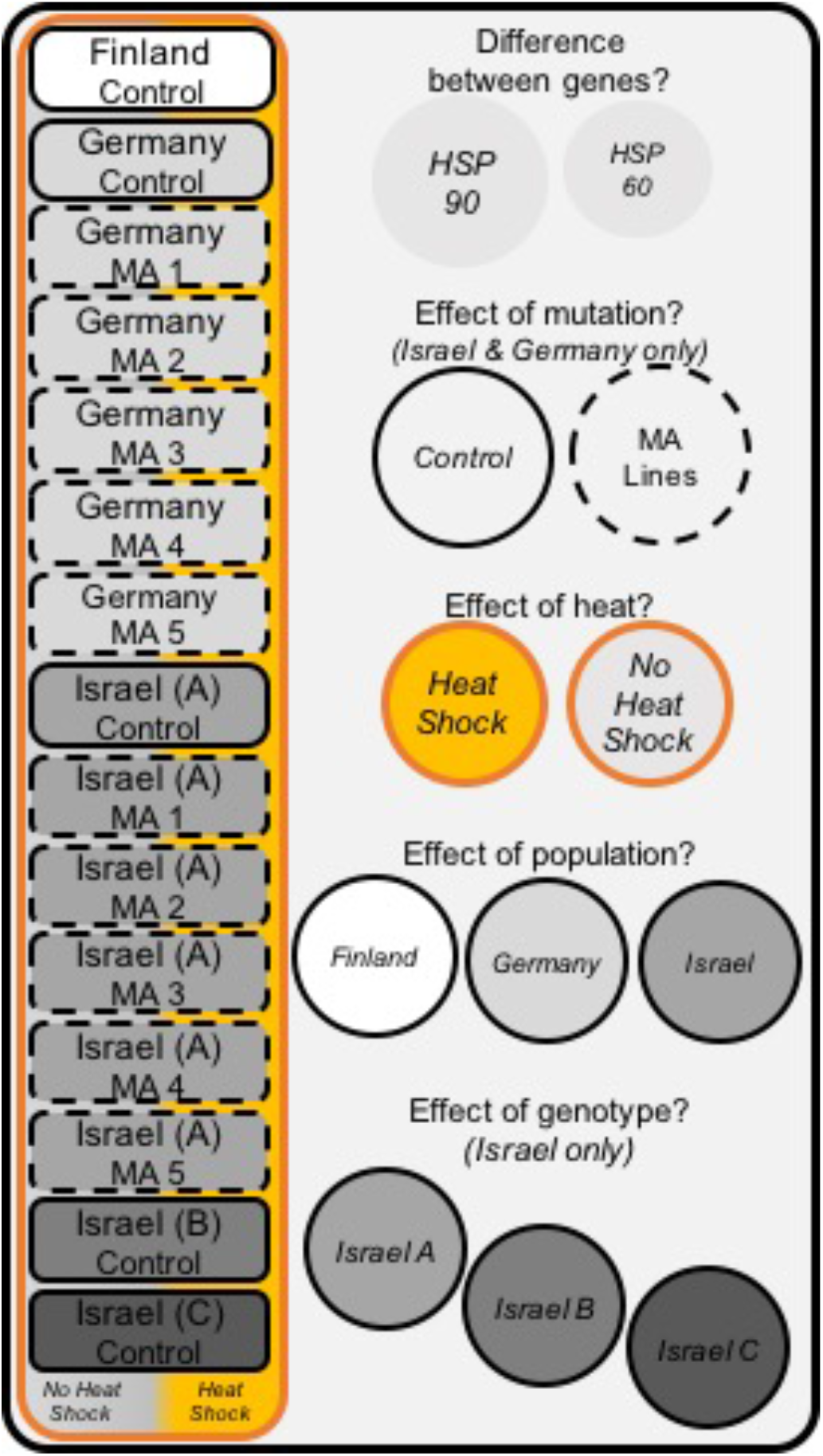
Experimental design showing all 15 genotypes assayed (rectangles on left) in triplicate to quantify HSP60 and HSP90 expression levels. Genotypes include descendants of original isolates from Finland, Germany, and three genotypes from Israel (solid border) and individuals from mutation accumulation lines derived from two of these genotypes (dashed borders). Assays were performed on individuals raised in a common laboratory environment exposed to one of two environmental conditions (no heat shock [No HS; gray background] or heat shock [HS; yellow background]). The five axes of comparison made possible using this design are summarized in the circles on the right.

### Heat Shock Exposure

To assess the effect of heat on HSP gene expression, replicate fourth generation *D. magna* from the same clutch were raised in pairs in 40 mL of ADaM in 50 mL plastic conical tubes at 18 °C. Two pairs of individuals per line were raised for each treatment (heat shock and non-heat shock control). After 15 days of growth and regular feeding, each individual was transferred to a 1.7 mL microcentrifuge tube. For each line, half of the individuals were placed in a 30 °C Corning LSE Digital Dry Bath inside of an 18 °C Percival incubator (heat shock), and the other half were placed in a Corning LSE Digital Dry Bath that was turned off and equilibrated to ambient temperature inside of the same 18 °C Percival incubator (no heat shock). Individuals were treated for 2 hours. After 2 hours, the media was removed and replaced with 300 μL 1X DNA/RNA Shield from the Zymo Research Quick-RNA Miniprep Kit. Samples were frozen immediately in liquid nitrogen and stored at −20 °C.

### RNA Extraction and Reverse Transcription

RNA was extracted from each sample independently using the Zymo Research Quick-RNA Miniprep kit according to the manufacturer’s protocol. Briefly, one *D. magna* individual in 1X DNA/RNA Shield was mixed with 300 μL RNA Lysis Buffer and ground with a microcentrifuge pestle. All centrifugations were done at 10,000 × g for 30 seconds unless specified with a Labnet Spectafuge 24D. After centrifugation through a DNA specific filter for 1 min, the flow-through was mixed with 600 μL ethanol, transferred to an RNA specific filter, and centrifuged. The bound RNA was then washed with 400 μL of RNA Wash Buffer and treated with a solution of 75 μL DNA Digestion Buffer and 5 μL DNase I for 15 min in order to destroy any remaining DNA. The digestion was centrifuged, and the remaining RNA was washed once with 400 μL RNA Prep Buffer and once with 700 μL RNA Wash Buffer. The final wash was done with 400 μL RNA Wash Buffer, and it was centrifuged for 2 min in order to remove any latent buffer. RNA was then eluted into a nuclease-free microcentrifuge tube with 50 μL DNase/RNase free water and stored at −20 °C. Concentration of RNA was measured using the Invitrogen Qubit RNA BR Assay with a Qubit 3.0 (Life Technologies). For each sample, 100 ng of total RNA per individual was reverse transcribed with random primers in a 20 μL reaction using the Promega GoTaq 2-Step RT-qPCR System according to the manufacturer's protocol. cDNA was then stored at −20 °C.

### Quantitative PCR

An RNA sequence for HSP60 was obtained from Steinberg et al. (2010) and the sequence for HSP90 from Kotov et al. (2006). Sequences were aligned to whole genome sequences of control lines from each population in this study using blastn (see Supplemental Data File A for alignments). Candidate control genes (succinate dehydrogenase, glyceraldehyde-3-phosphate dehydrogenase (GAPDH), and ubiquitin conjugating protein (UBC) for qPCR were selected from Heckmann et al. (2006).

Primers were designed using Primer3 to generate amplicons between 70 bp and 200 bp (Supplemental Table S0). After qPCR, the stability of each control gene was checked using RefFinder (Xie et al., 2012). Though UBC expression was previously observed to be somewhat responsive to heat in different *D. magna* populations (Jansen et al., 2017), we found it to be the most stable across treatments in our populations, so it was used as the control gene for this study. Primer efficiencies were assessed by serial dilution. Both target genes and UBC were found to have efficiencies of 100% (Supplemental Figure 1). Any primer pairs with estimated efficiencies slightly over 100% were assumed to have true efficiencies of 100%. Primer functionality and specificity were verified through end-point PCR using Qiagen Taq PCR Master Mix. Products were analyzed by gel electrophoresis. Amplicon lengths are as follows: HSP90 is 138 bp, HSP60 is 74 bp, and UBC is 90 bp.

qPCR was performed using the Promega GoTaq 2-Step RT-qPCR System according to the manufacturer’s protocol. Each 10 μL reaction included 5 μL GoTaq qPCR Master Mix, 2 μL each of 1 μM forward and reverse primers, and 1 μL of cDNA. Cycling conditions (CFX Connect, Bio-Rad) were 2 min at 95 °C for polymerase activation followed by 40 cycles of 15 sec of denaturation at 95 °C with 1 min at 55 °C of annealing and extension. Lastly, a melt curve from 55 °C to 95 °C was added at the end to verify no off-target amplification. Samples and genes were organized through the sample maximization method such that each plate only amplified one gene, but each plate had all samples (2-3 biological replicates per line and treatment). Three technical replicate reactions were performed on separate plates. Because each sample was represented in every plate, plates served as technical replicates (Derveaux et al., 2010).

### Data Analysis

In order to determine if any technical replicates were outliers, the mean of each sample x gene combination was calculated. Only replicates < 1 standard deviation from the mean (−1 < Z-score < 1) were included in the analysis. The relative quantity (RQ) of experimental genes (HSP90, HSP60) originally present in the sample was calculated using the mean C_q_ of the remaining replicates and the efficiency of the primer pair (E), normalized by the RQ of the reference gene (UBC) as described by Rieu and Powers (2009) to estimate normalized relative quantities (NRQ). NRQ values were log transformed prior to statistical analysis to correct for heterogeneity of variance (Rieu and Powers, 2009). The raw data can be found in Tables S6 and Table S7 for HSP90 and HSP60, respectively. Transformed data (using a log2(NRQ) transformation) are in Table S8 and Table S9, for HSP90 and HSP60, respectively.

We tested our log-transformed dataset for normality and homogeneity of variances. Using the Levene's test, the data for HSP90 (F13,70 = 1.56, p = 0.117) and HSP60 (F13,70 = 1.08, p = 0.388) suggest that there is homogeneity of variances. Through a Shapiro-Wilks test on the residuals of a multiple linear regression model including all data for both genes independently, HSP60 did not depart significantly from normality (W = 0.974, p = 0.0877) while HSP90 expression levels were found to have high non-normality (W = 0.811, p < 0.0001). As the data were already log_2_ transformed, there was no further transformation that improved the normality of the dataset. Q-Q plots of expression levels of both genes show a higher than predicted number of cases at both ends of the model (Supplemental Figure 2). However, because there is no non-parametric equivalent of a multi-way ANOVA, and ANOVA is robust to departures from normality (Knief and Forstmeier, 2020) such as those in this dataset, differences in means were tested using ANOVAs.

All ANOVAs were performed in RStudio. The full model tested the effects of gene (HSP60, HSP90), heat shock, mutation accumulation, population of origin, and genotype, and all interactions, on expression level using a 5-Way ANOVA (Model A in R code and Table S2). To test for mutation accumulation effects specific to HSP90 and HSP60, a model was made for each gene with all samples including both mutation accumulation lines and control lines with all populations using a 4-Way ANOVA (Models B and C respectively in R code and Tables S3 and S4). To test for population effects, in addition to Model A, two additional models, D and E, were made that included only control lines from each population (with all genotypes from Israel) for each gene using a 2-Way ANOVA (Tables S3 and S4). Lastly, two 2-Way ANOVA models were made using only Israel control lines for each gene to test if genotype has an effect on HSP90 or HSP60 expression (Models F and G in R code and Table S5). All models can be found in the supplemental tables and R code.

## Results

Our assay of gene expression levels for HSP60 and HSP90 allowed us to test for the effect of heat stress (30°C vs. 18°C), mutation accumulation (5 MA lines compared to control lines from both Israel and Germany), population effects (Israel, German, Finland) and genotype effects (three genotypes nested within the Israel population) in *D. magna* (Figure 1). Overall, HSP90 was expressed approximately 10-fold higher than HSP60 (F = 163.7, df = 1, p << 0.001; Table 1, Table S2, and Figure 2). This difference in expression was observed under both unstressed and heat shocked conditions (Figure 2).

**Figure 2.**
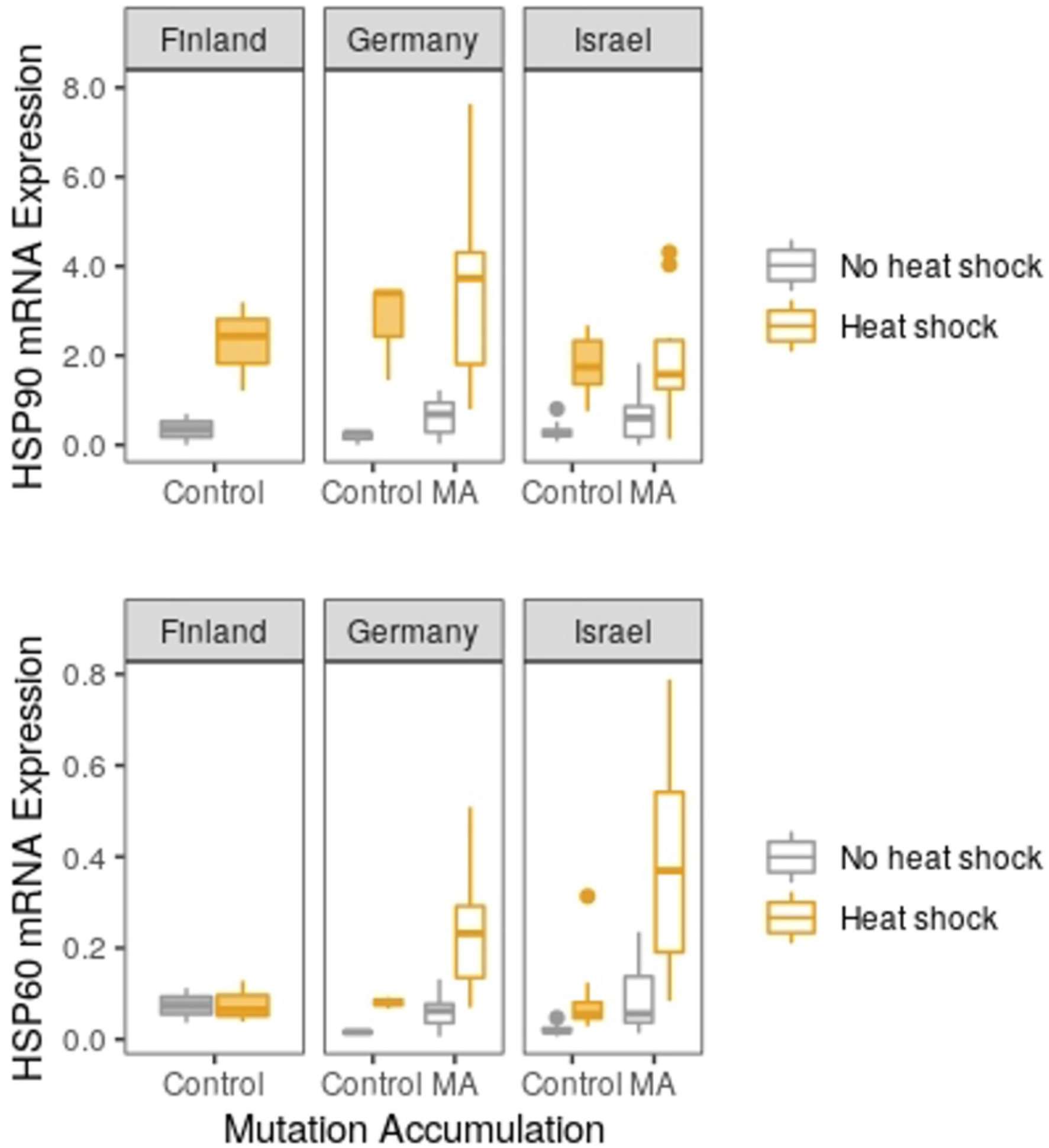
Gene expression for HSP90 (top) and HSP60 (bottom) in genotypes from three populations (Finland, Germany, and Israel) from individuals from mutation accumulation (unshaded) versus control lines (shaded) that were (yellow) and were not heat shocked (gray). Horizontal lines represent medians, boxes indicate quartiles and vertical lines illustrate the maximum value of 1.5 x IQR + the 75th percentile and the minimum value of the 25th percentile - 1.5 x IQR of the variance. Note: One outlier in Germany MA (HSP90 mRNA Expression = 12.64) was excluded from the graph of HSP90 expression to better visualize differences in medians; however, it is included in the ANOVA results in Table 1.

**Table 1.** Analysis of variance (ANOVA) for gene expression based on transcript abundance for HSP60 and HSP90 assayed in *Daphnia magna* originally collected from three populations (in Finland, Germany, and Israel), subject to mutation accumulation, and raised with and without heat shock. For complete ANOVA tables of all data partitions, see Supplemental Tables S2-S5; for the raw data used in this analysis, see Supplemental Tables S6 and S7.

| Data partitions | Factor | Df | Sum of Squares | F value | Pr(>F) |
| --- | --- | --- | --- | --- | --- |
| *All data* |  |  |  |  |  |
| *Main effects and 2/3-way interactions* | Population | 2 | 6.63 | 1.2611 | 0.2866 |
|  | **Gene** | 1 | 430.19 | 163.7158 | **< 0.0001** |
|  | **HeatShock** | 1 | 268.22 | 102.0753 | **< 0.0001** |
|  | **MutationAccumulation** | 1 | 41.24 | 15.6935 | **0.0001** |
|  | Population:Gene | 2 | 7.83 | 1.4898 | 0.2290 |
|  | Population:HeatShock | 2 | 3.63 | 0.6902 | 0.5032 |
|  | Gene:HeatShock | 1 | 7.65 | 2.9106 | 0.0902 |
|  | Population:MutationAccumulation | 1 | 1.73 | 0.6589 | 0.4183 |
|  | **Gene:MutationAccumulation** | 1 | 15.4 | 5.8607 | **0.0168** |
|  | HeatShock:MutationAccumulation | 1 | 1.37 | 0.5202 | 0.4720 |
|  | Population:Genotype | 2 | 7.37 | 1.4027 | 0.2494 |
|  | **Population:Gene:HeatShock** | 2 | 20.11 | 3.827 | **0.0241** |
| *HSP 60 only* |  |  |  |  |  |
| *Main effects* | Population | 2 | 0.819 | 0.3281 | 0.7214 |
|  | **HeatShock** | 1 | 92.641 | 74.2147 | **< 0.0001** |
|  | **MutationAccumulation** | 1 | 53.518 | 42.8737 | **< 0.0001** |
| *HSP 90 only* |  |  |  |  |  |
| *Main effects* | Population | 2 | 13.637 | 1.7017 | 0.1898 |
|  | **HeatShock** | 1 | 183.224 | 45.726 | **< 0.0001** |
|  | MutationAccumulation | 1 | 3.118 | 0.7782 | 0.3807 |
| *HSP 60 only, Israel only* |  |  |  |  |  |
| *Genotype Effects* | Genotype | 2 | 4.0676 | 3.0971 | 0.0823 |
|  | **HeatShock** | 1 | 16.6449 | 25.3471 | **0.0003** |
|  | Genotype:HeatShock | 2 | 3.9055 | 2.9737 | 0.0893 |
| *HSP 90 only, Israel only* |  |  |  |  |  |
| *Genotype Effects* | **Genotype** | 2 | 5.729 | 6.3804 | **0.0130** |
|  | **HeatShock** | 1 | 34.261 | 76.3196 | **< 0.0001** |
|  | Genotype:HeatShock | 2 | 1.048 | 1.1673 | 0.3442 |

Generally, heat shock increases the mean expression levels of both genes (F = 102.1, df = 1, p < 0.001; Table 1 and Figure 2), although the specific fold-change depends on the gene and population-of-origin (~6x increase, on average but in some cases as much as a 15x increase). Similarly, we observed higher expression levels for both genes in mutation accumulation lines relative to control lines (F = 15.7, df = 1, p < 0.0001; Table 1 and Figure 2), although the size of the increases were not as large as with heat shock (on average, 3.8x increase; Table 2). Individually, the effect of mutation accumulation was significant for HSP60 (F = 42.9, df = 1, p < 0.0001; Table 1 and Table S4), but not for HSP90 (F = 0.8, df = 1, p = 0.381; Table 1 and Table S3), though HSP90 expression levels were elevated in lines where mutations had accumulated, regardless of temperature (Table 2). There is evidence for a synergistic effect of heat and mutation accumulation, with much higher expression with the combination of both stresses (on average, 23x increase) than under either stress individually (Figure 2 and Table 2). Both factors, heat shock and mutation accumulation, tend to not only increase the mean expression levels, but also the variance in gene expression levels of both genes (Figure 2).

**Table 2.** Estimated mean expression levels for HSP60 and HSP90 assayed in *Daphnia magna* originally collected from three locations (Finland, Germany, and Israel), subject to mutation accumulation, and raised with and without heat shock. For Germany and Finland, one genotype each was sampled (GC and FC, respectively). For Israel, three individual genotypes were assayed (IA, IB, and IC). For complete ANOVA tables of all data partitions, see Supplemental Tables S2-S5; for the data used in this analysis, see Supplemental Tables S6 and S7.

| Gene | Population | Genotype | Mutation Accumulation | Heat Shock | Mean Expression (NRQ) Untransformed |
| --- | --- | --- | --- | --- | --- |
| HSP90 | Finland | FC | No | - | 0.350 |
|  |  | FC | No | + | 2.282 |
|  | Germany | GC | No | - | 0.181 |
|  |  | GC | No | + | 2.754 |
|  |  | GC | Yes | - | 0.674 |
|  |  | GC | Yes | + | 4.267 |
|  | Israel | IA, IB, IC | No | - | 0.314 |
|  |  | IA, IB, IC | No | + | 1.791 |
|  |  | IA | Yes | - | 0.636 |
|  |  | IA | Yes | + | 1.818 |
|  |  | IA | No | - | 0.124 |
|  |  | IA | No | + | 1.287 |
|  |  | IB | No | - | 0.392 |
|  |  | IB | No | + | 1.917 |
|  |  | IC | No | - | 0.425 |
|  |  | IC | No | + | 2.169 |
| HSP60 | Finland | FC | No | - | 0.074 |
|  |  | FC | No | + | 0.077 |
|  | Germany | GC | No | - | 0.015 |
|  |  | GC | No | + | 0.080 |
|  |  | GC | Yes | - | 0.059 |
|  |  | GC | Yes | + | 0.245 |
|  | Israel | IA, IB, IC | No | - | 0.021 |
|  |  | IA, IB, IC | No | + | 0.090 |
|  |  | IA | Yes | - | 0.090 |
|  |  | IA | Yes | + | 0.384 |
|  |  | IA | No | - | 0.010 |
|  |  | IA | No | + | 0.135 |
|  |  | IB | No | - | 0.019 |
|  |  | IB | No | + | 0.042 |
|  |  | IC | No | - | 0.035 |
|  |  | IC | No | + | 0.094 |

In terms of intraspecific variation in gene expression, levels did not vary based on which population the genotypes originated from (Finland, Germany, and Israel; F = 1.26, df = 2, p = 0.29; Table 1), although there was one interaction effect observed (population x gene x heat shock; Table 1). This was driven by the fact that there was an effect of heat shock in all three populations for HSP90, but only for two of the three populations for HSP60 (not Finland; see Table S2 for post-hoc pairwise contrasts). Surprisingly, there is a genotype effect for HSP90 expression levels (comparing genotypes IA, IB, and IC from Israel, excluding all MA lines; F = 6.4, df = 2, p = 0.01; Tables 1 and 2 and Figure 3), but no significant genotype effect was observed in HSP60 (F = 3.1 df = 2, p = 0.08; Table S5).

**Figure 3.**
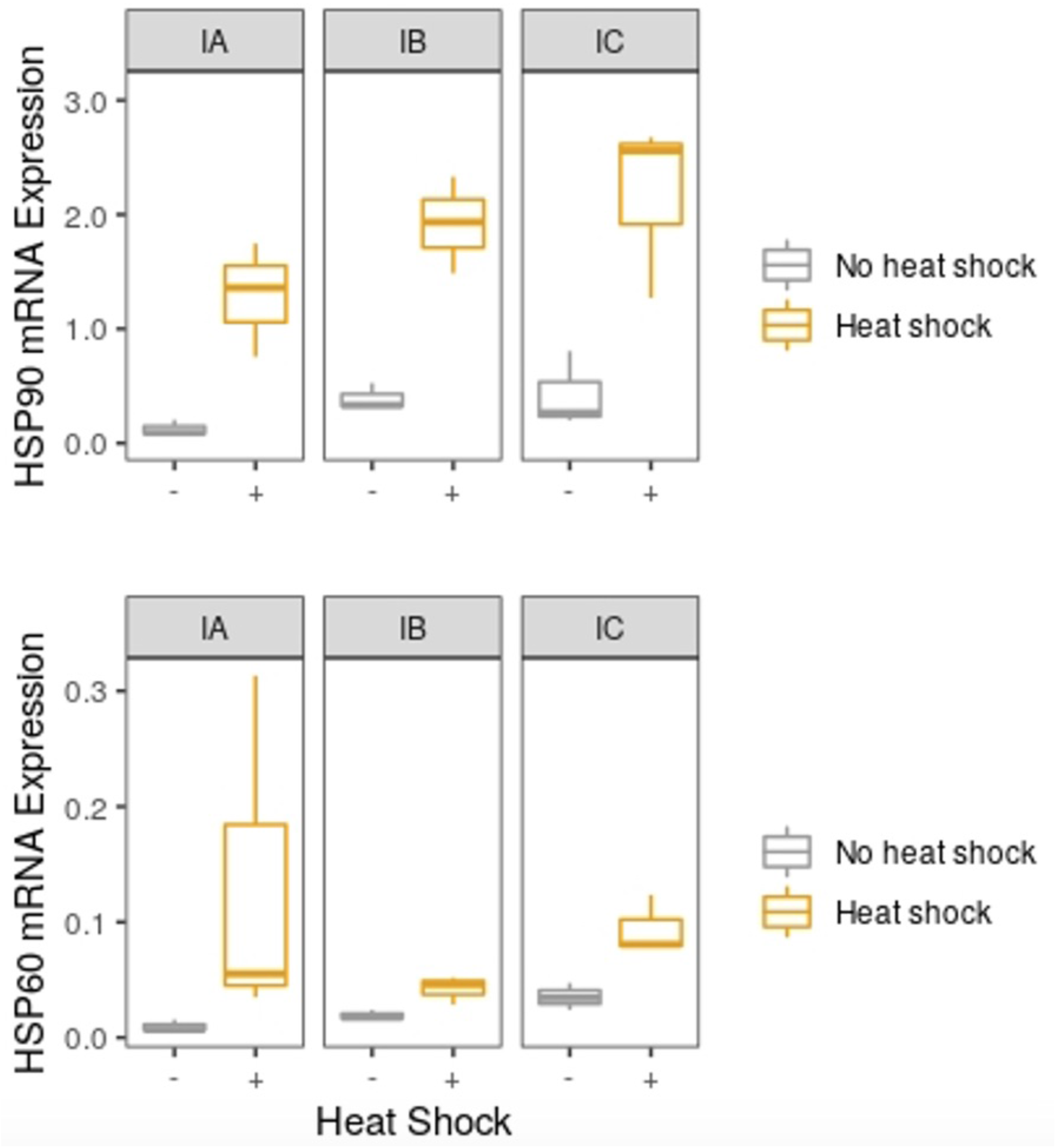
Gene expression levels for HSP90 (top) and HSP60 (bottom) with exposure to heat shock (yellow) and without heat shock (gray) for three genotypes from Israel (data for ANOVAs appears in Table S5). Horizontal lines represent medians, boxes indicate quartiles and vertical lines illustrate the maximum value of 1.5 x IQR + the 75th percentile and the minimum value of the 25th percentile - 1.5 x IQR of the variance.

## Discussion

The HSP genes are members of a large and diverse family and play a variety of important roles in responding to extrinsic and intrinsic sources of stress (Neuhaus-Steinmetz et al., 1997; Kim et al., 2014; Liu et al., 2015). Here, we performed a controlled laboratory experiment to compare the expression profiles of HSP90 and HSP60 with and without heat stress and mutation accumulation, and compare expression levels and changes among populations and genotypes collected along a latitudinal gradient. While HSP90 has long been referred to as a mutational “capacitor” because of its major role in protein folding and the large number of proteins it interacts with (Schopf et al., 2017), the role of HSP60 in the stress response is less well understood given its localization primarily to the mitochondria (Magnoni et al., 2014). Recent studies have reported the highest direct estimates of spontaneous mutation rates in animals from mutation accumulation experiments with *D. magna* (Ho et al., 2019, Ho et al., 2020). Their importance as an ecological and environmental model system make an understanding of their stress response and their ability to buffer the phenotypic effects of mutation of particular interest (Latta et al. 2015).

Overall, we find that HSP90 is expressed ~10x more than HSP60 in *D. magna* (Table 1 and Figure 2). This corroborates previous work that shows HSP90 constitutes approximately 1-2% of the total protein content of eukaryotic cells (Borkovich et al., 1989) and, in yeast, is known to interact with up to 20% of proteins (Taipale et al., 2010). As expected, we found both genes have a robust heat shock response in terms of HSP 90 and HSP60 expression increases (Table 1 and Figure 2). Heat shock destabilizes folded proteins, and elevated HSP expression protect against exposure of hydrophobic segments, aggregation, and misfolding (Kimura et al., 1993, Vabulas et al., 2010). It is known that HSP60 is upregulated in response to heat (Martin et al., 1992) and oxidative stress in *D. melanogaster* (Singh et al., 2009), but a multi-faceted, rapid HSR may be especially important for aquatic animals living in shallow water because they can experience major temperature fluctuations (Feder and Hoffman, 1999). We also observed an increase in gene expression in mutation accumulation lines relative to controls, especially in HSP60 (Table 1). That this response is especially acute in HSP60 may be related to the higher mutation rates observed in the mtDNA genome, relative to the nuclear genome, in animals (although mtDNA mutation rates are notoriously difficult to accurately measure [Schaack et al., 2020]). The greater upregulation of HSP60 in response to mutation accumulation underscores the importance of examining the potential of other HSPs (in addition to HSP90) as potential mutational capacitors (Rutherford and Lindquist, 1998).

In addition to looking at the effects of heat shock and mutation, we were also interested in differences within and among populations in both their baseline levels of expression and their response to stress. Surprisingly, there are no significant differences in gene expression at either locus among populations (Table 1), despite the major abiotic differences between these locales in mean annual temperatures (approximately 2, 10, and 21 degrees C in Finland, Germany, and Israel, respectively; Rohde and Hausfather 2020). It could be that the evolution of HSRs depend more on maximum temperatures or temperature fluctuations, which exhibit a much smaller range of only ~10 and 7 degrees, respectively (Table S1; Hofmann and Somero, 1996; Gehring and Wehner; 1995; Cambronero et al., 2018). However, given that the genotypes used in this study have extremely high identity in the coding regions of these loci (>99% of sites are identical [418/422] for HSP60 and 735/741 for HSP90; Supplemental Data Files), differences in gene expression are more likely due to variation at promoter regions or other loci in the genome which may serve to regulate HSP expression. While our predictions about population differences did not bear out, there is a difference in expression among genotypes within a population (IA, IB, and IC genotypes from Israel) for HSP90 and a non-significant trend for HSP60. Interestingly, the genotype with the highest levels of heat-induced gene expression (Figure 3) is also the genotype with the highest mtDNA base substitution mutation rate among those from Israel (Ho et al. 2020), further supporting the notion that HSP expression could provide a buffer against high mutation rates.

Our study provides strong evidence for the synergistic effects of multiple stresses on HSP expression. In all cases where a given genotype was assayed with and without heat shock and mutation accumulation, the combination of the two stressors resulted in an increase in the expression levels that was an order of magnitude greater than the increase observed when only one stress is applied (Table 2). Furthermore, the variance was greater in the cases where two stressors were applied (Figure 2). This has important implications for *Daphnia,* and other species, as global climate change does not only lead to different mean and maximum temperatures and temperature fluctuations. Changing climate can alter exposure to UV or other atmospheric mutagens, and can also reduce the availability of freshwater aquatic habitats caused by drought or sea level rise. Reduced habitat availability will likely reduce effective population sizes for species like *D. magna,* and thus a further increase in their already high mutation rates (reviewed in Lynch et al. 2016; Ho et al., 2020). If HSPs can buffer against not only thermal stress but the accumulation of mutations, they could enable the evolution of higher mutation rates. While spontaneous mutations are known, to be, on average deleterious, beneficial mutations do occur. Ultimately, increases in genetic variation provide the evolutionary escape hatch or opportunity for rapid adaptation (Swings et al., 2017) necessary to tolerate or thrive in increasingly stressful environments.

## Acknowledgements

We would like to thank Theresa Steele and Aziz Ouedraogo for their invaluable assistance during the assay. We would like to thank Dieter Ebert for supplying the animals used to initiate the mutation accumulation experiment. We thank Kelly McConville for her statistical advice. We would also like to acknowledge our funding sources: awards from Reed College to HS and from National Institute of General Medical Sciences of the National Institutes of Health (GM132861) and National Science Foundation (MCB-1150213) to SS.

## Author Contributions

HS conceived of the study, carried out lab work, performed data analyses, and collaboratively wrote and edited the manuscript; JC assisted with experimental design, helped with lab work, data analysis, providing crucial comments and input to develop and improve the manuscript; EKHH extracted sequence data for primer design and analysis and performed alignments; SS supplied the lineages used in the study and helped/supervised the experimental design, live animal exposures, molecular assays, data analysis and interpretation, and writing and editing manuscript. All authors gave final approval for publication and agree to be held accountable for the work performed therein.

## Data Availability Statement

All raw and transformed data used in this study are in Supplemental Tables S6, S7, S8, and S9. All R code (SuppFile1) and sequence data (SuppFile2) have been uploaded as a Supplemental Files.

## Supplemental Materials

### Supplemental Data Files

SuppFile1_All_R_Code.txt

SuppFile2_Zipped_SequenceData.zip

### Supplemental Tables

Supplemental Tables S0-S9 are available, each on a single sheet, in a .xls workbook (SupplementalTables_122720.xlsx).

**Table S0.** Primers used for RT-qPCR with *Daphnia magna* cDNA extracts in this study.

**Table S1a, b, c.** Sample collection data for *Daphnia magna* and sample sizes for in this study.

**Table S2.** Complete multifactor ANOVA table conducted using the entire dataset of expression level data for *Daphnia magna* across genotypes, heat shock treatments, with and without mutation accumulation, including main and interaction effects. All R code is available in a supplemental file.

**Table S3.** Partitioned ANOVAs performed using HSP90 expression levels based on log2(NRQ) data found in Table S8.

**Table S4.** Partitioned ANOVAs performed using HSP60 expression levels based on log2(NRQ) data found in Table S9.

**Table S5.** Partitioned ANOVAs performed using HSP90 and HSP60 expression levels based on log2(NRQ) data for only Israel lines to test for genotype effects.

**Table S6.** HSP90 expression levels in *Daphnia magna* before log transformation (used in Figure 2 and Figure 3).

**Table S7.** HSP60 expression levels in *Daphnia magna* before log transformation (used in Figure 2 and Figure 3).

**Table S8.** Calculations of the log2(NRQ) for HSP90 across all available genotypes and biological replicates used for all statistics unless otherwise specified.

**Table S9.** Calculations of the log2(NRQ) for HSP60 across all available genotypes and biological replicates used for all statistics unless otherwise specified.

### Supplemental Figures

(see below, and SuppFig1.jpg and SuppFig2.jpg)

### Supplemental Figures

**Supplemental Figure 1:**
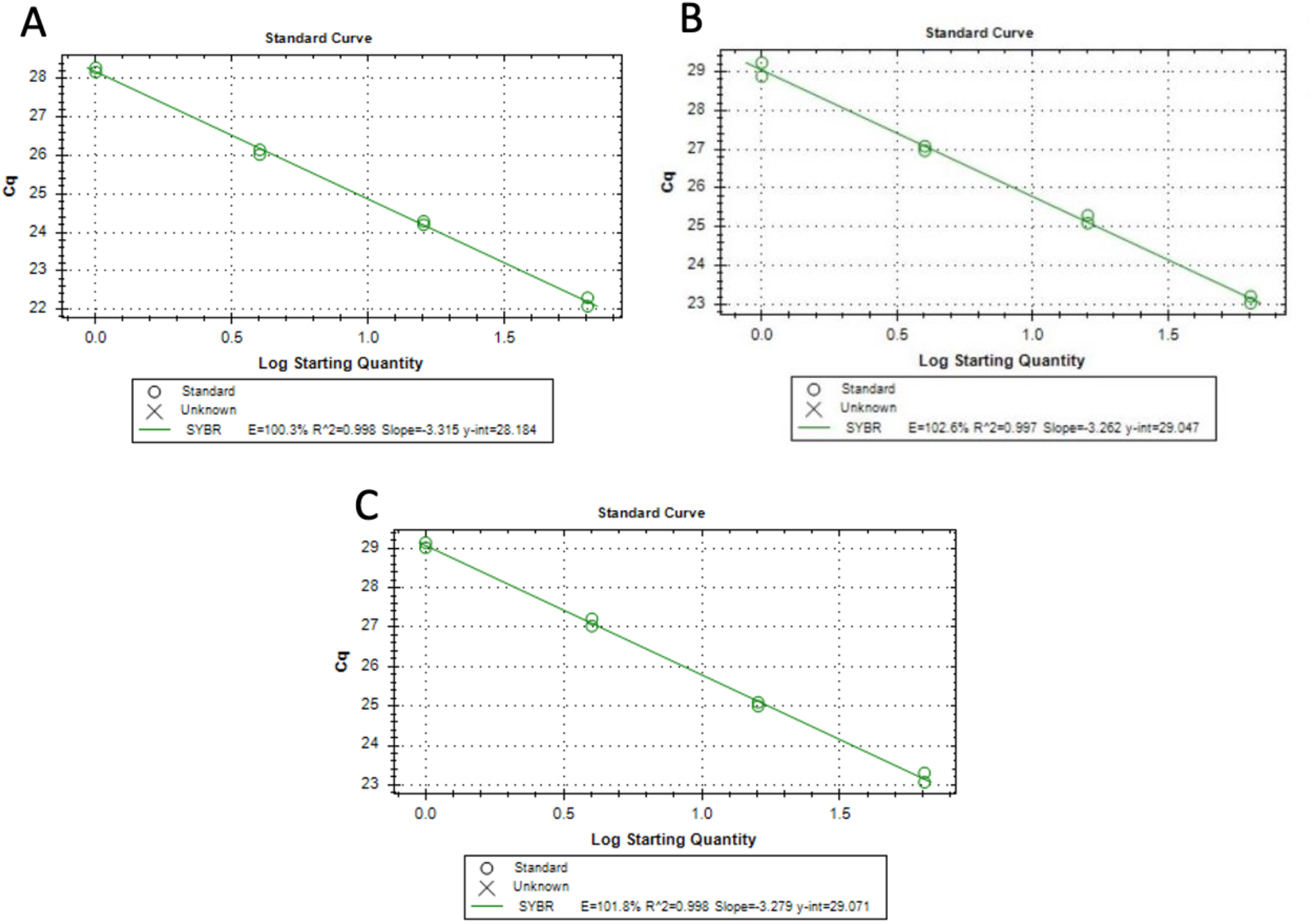
Amplification Curves of Dilution Series for qPCR Primers. Each standard curve was made by using the standard qPCR reaction mix and thermocycler program with two replicates of a dilution series of 1, 1/4, 1/16, and 1/64 of the original cDNA concentration. A) Standard amplification curve of HSP90 with an efficiency = 100.3%, B) standard amplification curve of HSP60 with efficiency = 102.6%, C) standard amplification curve of UBC with efficiency = 101.8%.

**Supplemental Figure 2.**
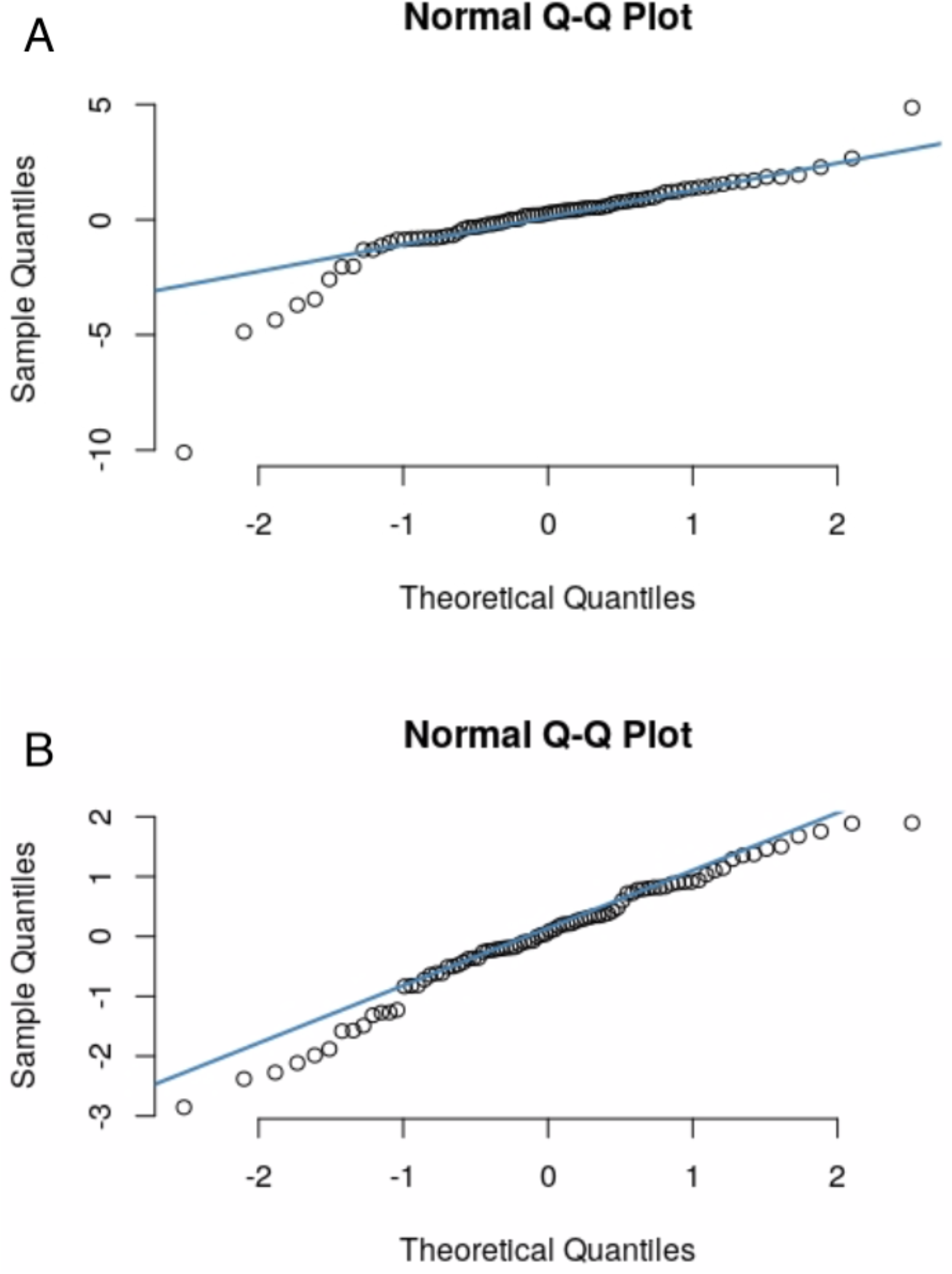
Q-Q plots of HSP90 mRNA expression levels. **(A) and HSP60 mRNA expression levels (B).** Q-Q plots were made from residuals of a multiple linear regression model using all samples for both genes independently.

